# Coupling between cerebral blood flow and cerebral blood volume: Contributions of different vascular compartments

**DOI:** 10.1101/202838

**Authors:** Roman Wesolowski, Nicholas P. Blockley, Ian D. Driver, Susan T. Francis, Penny A. Gowland

## Abstract

A better understanding of the coupling between changes in cerebral blood flow (CBF) and cerebral blood volume (CBV) is vital for furthering our understanding of the BOLD response. The aim of this study was to measure CBF-CBV coupling in different vascular compartments during neural activation. Three haemodynamic parameters were measured during a visual stimulus. Look-Locker Flow-sensitive Alternating Inversion Recovery (LL-FAIR) was used to measure changes in CBF and arterial CBV (CBV_a_) using sequence parameters optimised for each contrast. Changes in total CBV (CBV_tot_) were measured using a Gadolinium based contrast agent technique. Haemodynamic changes were extracted from a region of interest based on voxels that were activated in the CBF experiments. The CBF-CBV_tot_ coupling constant α_tot_ was measured as 0.16±0.14 and the CBF-CBV_a_ coupling constant α_a_ was measured as 0.65±0.24. Using a two compartment model of the vasculature (arterial and venous), the change in venous CBV (CBV_v_) was predicted for an assumed value of baseline arterial and venous blood volume. These results will enhance the accuracy and reliability of applications that rely on models of the BOLD response, such as calibrated BOLD.

## Introduction

The relationship between changes in cerebral blood flow (CBF) and cerebral blood volume (CBV) is critical for an accurate understanding of the haemodynamics that underlie blood oxygen level dependent (BOLD) fMRI, and a greater understanding of cerebral haemodynamics will lead to more precise quantification of the BOLD response [1,2]. Early studies suggested that the majority of CBV change in response to neuronal activation occurs in venous vessels [3,4]. This led to the adoption of a CBF-CBV coupling model based on a power law relationship that was characterised using PET measurements of CBF and total CBV (CBV_tot_) [5]. However, it is now well known that arterial CBV (CBV_a_) also increases on activation and to a much greater degree than total CBV, despite the lower baseline CBV of the arterial compartment [6–8]. Therefore, the use of a CBF-CBV_tot_ coupling constant will overestimate changes in venous CBV. This led to measurements of a CBF-CBV_v_ coupling constant using the CBV_v_ sensitive VERVE technique [9,10]. Despite this, the coupling between CBF and CBV_a_ is still poorly understood in humans, with the majority of research having been performed in rats [6,7]. Whilst changes in CBV_a_ are typically invisible in the context of standard BOLD fMRI, they become significant in studies utilising intravascular contrast agents [11] or hypoxic hypoxia [12], where an arterial signal change occurs due to the presence of paramagnetic contrast agent or deoxyhaemoglobin, respectively, in the arterial blood volume, the latter having implications for the application of these methods in cerebrovascular disease where patients may have a reduced arterial oxygen saturation.

One reason why there are only a small number of published studies examining the relationship between these parameters is the limited number of techniques for measuring CBV_tot_, CBV_a_ and CBV_v_ in humans. Fractional changes in CBV_tot_(ΔCBV_tot_) during a stimulus have been measured via an infusion of a Gadolinium based contrast agent [13–15]. Such experiments rely on measuring changes in the stimulus evoked BOLD response as a function of intravascular contrast agent concentration, which can either be increased using an extended infusion, or decreased by clearance of a bolus of contrast agent via the kidneys [16]. Vascular Space Occupancy (VASO) has also been shown to provide a method for the assessment of total CBV [17]. However, it is not possible to measure ΔCBV_tot_ using VASO without prior knowledge of the baseline CBV_tot_, therefore, making this technique unsuitable for the study of CBF-CBV_tot_ coupling. More recently, arterial spin labelling (ASL) based approaches have emerged for the measurement of CBVa: Inflow-based VASO (iVASO) [18] and Look-Locker Flow Alternating Inversion Recovery (LL-FAIR) [8]. LL-FAIR combines FAIR ASL technique with Look-Locker echo planar imaging (LL-EPI) sampling [19] to sensitise the signal to either CBF [20] or CBV_a_ [8] depending on the sequence parameters. This technique is also presented in the literature as ITS-FAIR [21] and QUASAR [22]. Since the LL-FAIR technique is capable of measuring transit time it is able to account for changes in transit time that may occur during neuronal activation. LL-FAIR also has a higher signal to noise ratio (SNR) per unit time for measurement of perfusion than conventional FAIR [21]. Finally, measurements of CBV_v_ have been performed using the Venous Refocusing for Volume Estimation (VERVE) technique [23] or hyperoxia BOLD contrast [24] methods. However, VERVE is hampered by assumptions regarding oxygenation changes during activation and hyperoxia BOLD contrast by relatively low SNR.

In this study LL-FAIR based measurements of CBF and CBV_a_ are acquired alongside estimates of ΔCBV_tot_ using a Gadolinium based contrast agent technique. The relationship between the resultant haemodynamic parameters is assessed using a power law relationship, building upon early studies of CBF-CBV coupling [5], the calibrated BOLD method [1,2] and BOLD modelling studies [3,4]. This analysis yields CBF-CBV_a_ and CBF-CBV_tot_ coupling constants. These measurements are also used to predict the change in CBV_v_ for an assumed arterial and venous volume fraction.

## Methods

### Imaging

This study was approved by the University of Nottingham Medical School Ethics Committee. Eight healthy volunteers aged 20 to 31 years (mean 24± 3, mean ± standard deviation) gave written consent and were scanned as part of this study. A schematic diagram of the experimental protocol is given in Fig. 1a. Data were acquired on a Philips Achieva 3 T system (Philips Healthcare, Best, Netherlands), using a body transmit coil and 8 channel SENSE head receive coil. All three haemodynamic parameter measurements were acquired with a common resolution of 3 × 3 × 5 mm^3^, matrix size of 64 × 64 and SENSE factor 2. Measurements of CBF and CBV_a_ were limited to a single slice acquisition at the time these experiments were performed. Therefore, an initial functional localiser scan was performed in order to select a single axial slice through the visual cortex with the largest region of BOLD activation. This slice prescription was then used throughout the rest of the experiment.

**Figure 1.**
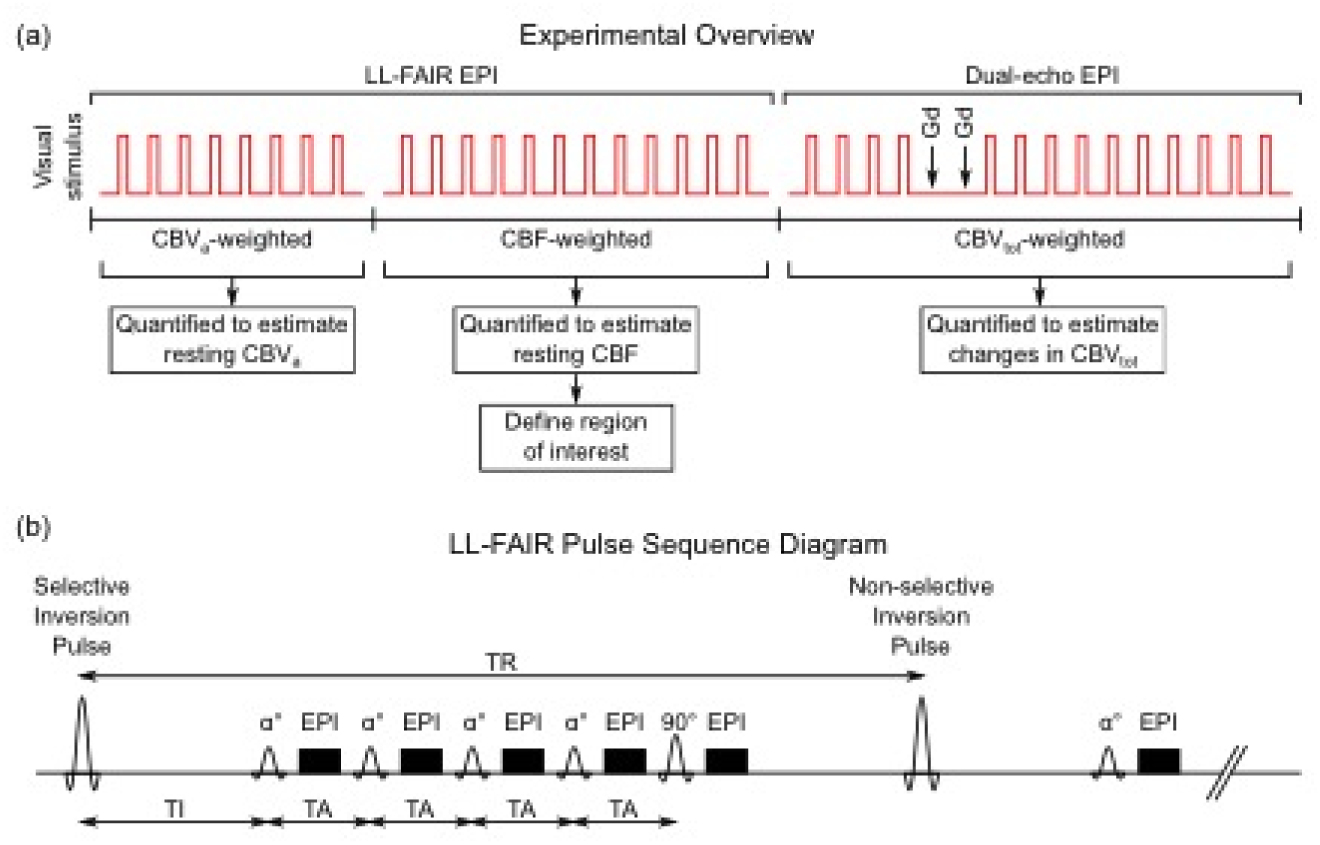
(a) Overview of the experimental protocol including visual stimulus blocks (red lines), contrast agent injections (arrow Gd), experimental techniques and processing steps. (b) Example pulse sequence diagram of LL-FAIR showing five readouts. The final readout pulse had a flip angle of 90°, preceeded by four lower flip angle α^0^ pulses. The sequence is repeated with repetition time (TR), the first readout pulse occurs following an initial inversion delay (TI) and readouts are separated by a time interval (TA).

Visual stimulation was provided by red LED goggles flashing at 8 Hz. Lights were ON 19.2 s and OFF for the remainder of the 60 s cycle. The number of stimulus cycles varied reflecting the differing SNR of each method: 8 cycles were collected for CBV_a_ measurements, 12 cycles for CBF (which has lower CNR) and 14 cycles for CBV_tot_.

A LL-FAIR acquisition was used for both CBF and CBV_a_ measurements. The combination of a Look-Locker acquisition with a FAIR preparation enables the tagged bolus to be tracked through the macrovascular and into the tissue. The signal can be sensitized to CBVa or CBF by using the appropriate combination of sequence parameters, and a kinetic model can be fit to quantify these parameters. In both cases the thickness of the inversion slab was alternated between 30 mm and 200 mm for label and control conditions, respectively. Fig. 1b displays the main features of the LL-FAIR pulse sequence. The sequence parameters for the CBF measurement were an initial inversion delay TI = 600 ms, time interval between EPI readouts TA = 350 ms (resulting in an inversion time range of 600 ms to 2000 ms), flip angle θ = 35° and 5 readout pulses with vascular crushing (bipolar lobe of 5 ms duration per lobe and amplitude of 15 mT). For the CBV_a_ measurement, sequence parameters were: TI = 150 ms, TA = 100 ms (resulting in an inversion time range of 150 ms to 1950 ms), θ = 50°, with 19 readout pulses. In both CBF and CBV_a_ measurements the shortest achievable echo time of 16 ms was used and the final LL-FAIR pulse had a flip angle of 90° to maximise SNR. The LL-FAIR scheme was performed with in-plane pre- and post-saturation pulses to provide signal suppression of the imaging slice, thus reduce any offset signals due to imperfections between the selective and non-selective RF inversion pulses. The application of a 90° pulse at the end of each TR simplified the modelling as it ensured each tag/control acquisition is independent, removing the need for an iterative fit of the data to be performed. The TR between inversion pulses was 2.4 s, resulting in a label/control pair being collected every 4.8 s.

For the measurement of CBV_tot_, dual-echo GE-EPI images were acquired with TE = 13/35 ms, TR = 1.2 s and 3 slices. Two single doses (0.2 ml kg^−1^) of Gadoteridol (ProHance, Bracco Imaging SpA, Milan, Italy) were injected, the first bolus at the beginning of the 5^th^ stimulus cycle and the second at the beginning of the 6^th^ cycle. For cycles 5 and 6, the visual stimulus was not presented and data from these cycles was not used in the estimation of ΔCBV_tot_. The final 10 cycles of the visual stimulus, post contrast agent injection, were acquired at different contrast agent concentration levels during clearance of the contrast by the kidneys.

### Analysis

For each subject, CBF, CBV_a_ and CBV_tot_ data sets were first realigned within each data set and then across all data sets using SPM5 (5 mm FWHM Gaussian smoothing kernel and second Degree B-Spline interpolation) [25]. For the CBF and CBV_a_ data, the images from the final LL-FAIR readout pulse (90° flip angle) of each volume acquisition (TR period) were realigned, and the transforms then applied to the images acquired from the other LL readout pulses within the corresponding TR period. Difference images (computed from the subtraction of consecutive label and control pairs) were calculated to give time series of CBF- and CBV_a_-weighted images. Average CBF and CBV_a_-weighted time series during the visual stimulus cycle were then formed by averaging across cycles, accounting for jittering in the data relative to the stimulus paradigm. The CBV_tot_ datasets were realigned using the images acquired at the first echo, and the motion transforms were then applied to the second echo data. Data for each echo time was then down-sampled to produce a complete time series with a 2.4 s temporal resolution (matching the CBF and CBV_a_ datasets), and only a single slice co-registered to the CBF and CBV_a_ datasets was retained. The time series of R_2_^*^ values was then calculated from this realigned, down-sampled, dual echo data.

CBV_a_-weighted difference images were quantified using a two-parameter fit, as described in [8], to measure changes in arterial transit time and CBV_a_. CBF-weighted difference images were initially analysed using a two-parameter fit for capillary transit time and CBF [20]. However, since the data had lower SNR than the CBV_a_ data, a two-parameter fit for each time point was found to increase the noise in CBF measures. Therefore, a mean estimate of the transit time at baseline and on activation was computed, and used in a one-parameter model fit. The fractional change (absolute change in the relevant parameter with respect to its absolute baseline value) in CBF (ΔCBF) and CBV_a_ (ΔCBV_a_) in response to the visual stimulus was then calculated from quantified CBF and CBV_a_ maps. The fractional change in CBV_tot_ (ΔCBV_tot_) was calculated by considering the effect of the contrast agent on the R_2_^*^ (transverse relaxation rate) changes that occur during the BOLD response, as shown previously by using an infusion to gradually increase the contrast agent concentration [15]. However, in this study two bolus injections of contrast agent were used to raise the initial blood contrast agent concentration, which gradually decreased due to washout through the kidneys.

In the analysis, four stimulus cycles prior to the contrast agent injections provided a baseline. Stimulus cycles following immediately after the injections were discarded to allow for recirculation of the contrast agent, leaving the final 9 cycles for analysis. In all other aspects, this method is the same as in previous reports and resulted in an estimate of ΔCBV_tot_ across the activation cycle [11,15].

Activated regions were generated for each subject using a correlation analysis applied to the quantified CBF data; regions of interest (ROI) were defined based on CBF statistical maps with a threshold p<0.01 (uncorrected).

### Estimation of CBF-CBV coupling

The coupling between CBF and CBV during neuronal activation was calculated assuming the power law relationship between CBF and CBV_tot_,

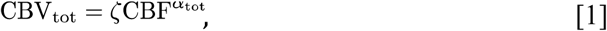

where ζ and α_tot_ were previously measured as 0.80 and 0.38, respectively, in rhesus monkeys using a hyper/hypocapnia challenge [5]. Considering the fractional change in CBV_tot_ (ΔCBV_tot_) and CBF (ΔCBF), Eq. [1] can be rearranged to calculate the CBF-CBV_tot_ coupling constant α_tot_ (Grubb’s constant):

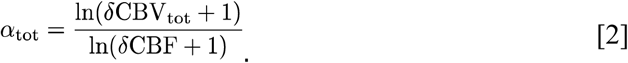

We assumed a similar power law relationship, α_a_, between ΔCBF and the fractional changes in CBV_a_ (ΔCBV_a_):1

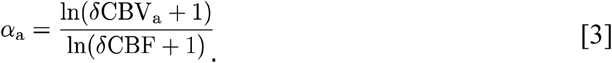

Using these equations, Grubb’s constant, α_tot_, and the CBF-CBV_a_ coupling constant α_a_ were estimated on a per subject basis. Mean steady state fractional changes in these parameters were extracted from the common ROI for each subject within active (ON) and baseline (OFF) time windows defined to be 9.6 – 19.2 s and 40.8 – 60 s, respectively, across the averaged stimulus cycle (Fig. 2). Estimates of α_tot_ and α_a_ were then computed using Eq. [2] and [3], respectively.

**Figure 2.**
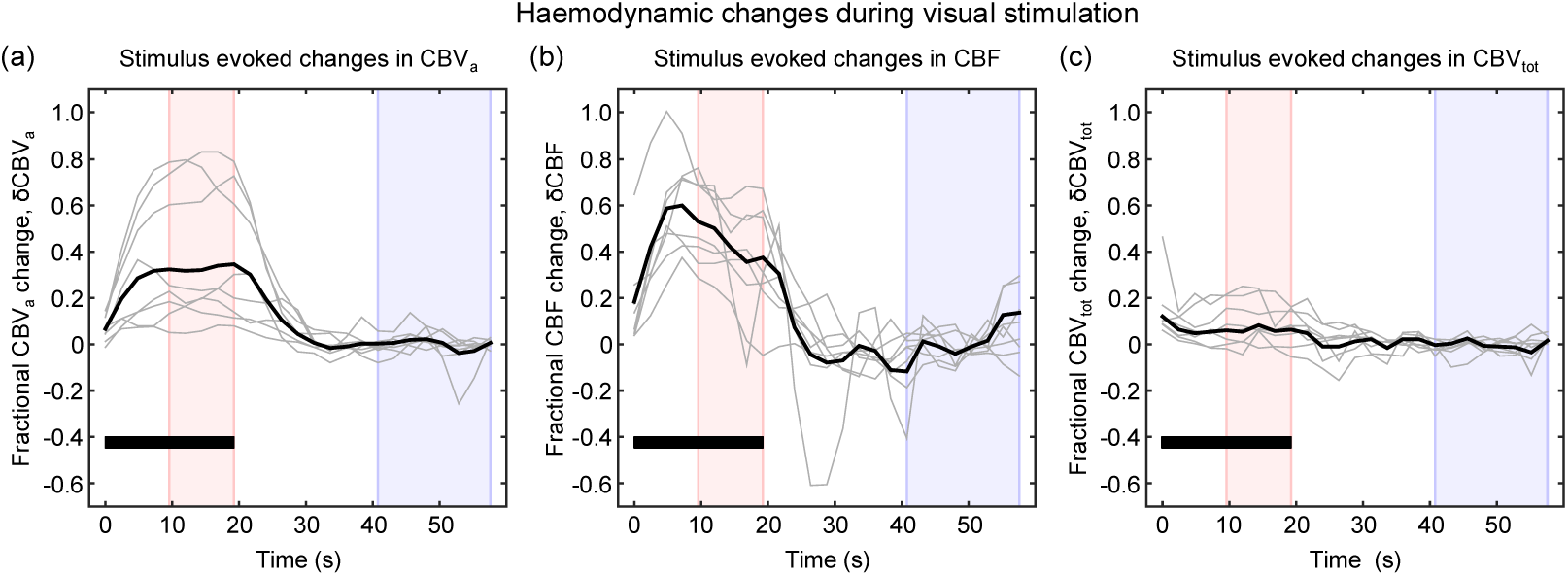
Haemodynamic changes during visual stimulation: (a) fractional change in arterial cerebral blood volume (δCBV_a_), (b) fractional change in cerebral blood flow (δCBF) and (c) fractional change in total cerebral blood volume (δCBV_tot_). Timecourses displayed for all subjects (grey lines) and group mean (black solid line). The visual stimulus period is denoted by a solid black bar and averaging windows highlighted for ON (pink) and OFF (blue) conditions.

In the absence of direct measurements of CBV_v_, we estimated the fractional change in venous CBV (ΔCBV_v_) using a simple model of the vascular compartments. Changes in total blood volume were approximated as a volume-weighted sum of two compartments: arterial and venous,

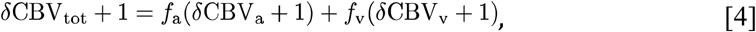

where *f*_a_ and *f*_v_ are the volume fractions assigned to arterial and venous blood volume compartments (*f*_a_=CBV_a_/CBV_tot_, *f*_v_=CBV_v_/CBV_tot_), respectively, and here it is assumed that *f*_a_ + *f*_v_ =1. The capillary volume is assumed to be distributed between the arterial and venous compartments. By rearranging Eq. [4], ΔCBV_v_ can be predicted as a function of arterial and venous volume fractions *f*_a_ and *f*_v_:

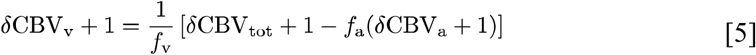

## Results

Figure 2 displays the time courses for the fractional change in each haemodynamic measure (CBV_a_, CBF and CBV_tot_), per subject (grey lines), and the group mean weighted by the number of voxels in each subject’s CBF-derived ROI (solid black line) (see Table 1). Table 1 lists the fractional changes measured for each haemodynamic parameter, along with the predicted value of ΔCBV_v_ estimated assuming *f*_*a*_=0.3 (*f*_*v*_=0.7), this choice of value is discussed below. Measurements of ΔCBV_tot_ and δCBV_a_ were found to be statistically significantly different from zero across the group (one-sample t-test, p<0.05), with estimates of α_tot_ and α_a_ provided in Table 1 for each subject. In contrast, the estimated value of δCBV_v_ was not statistically significantly different from zero (p=0.33), which would suggest a CBF-CBV_v_ coupling constant, α_v_, of close to zero. Group means and standard deviations in Table 1 are weighted by the number of voxels in each subject’s ROI.

**Table 1.**
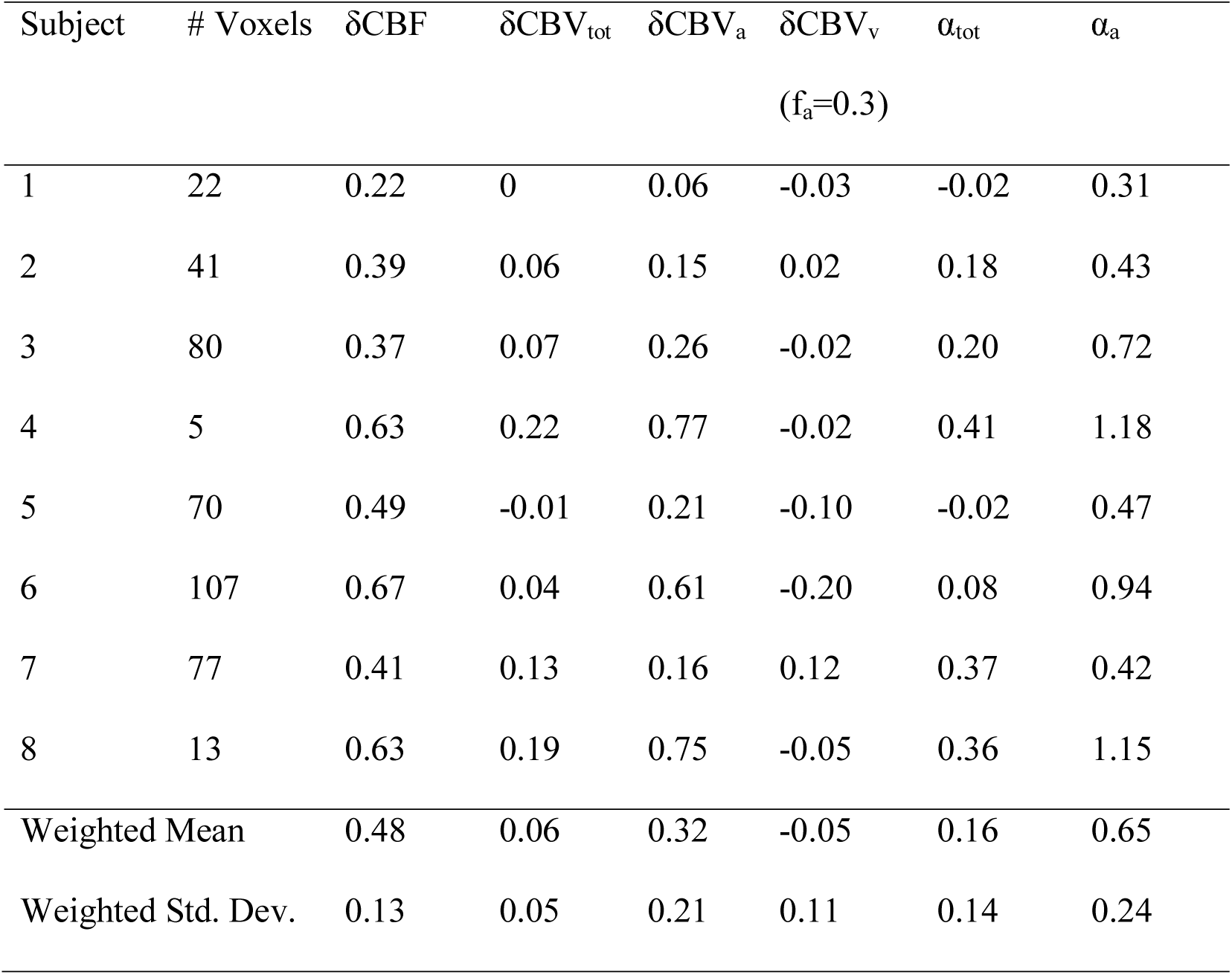
Experimental measurements of steady state fractional changes in cerebral blood flow (δCBF), arterial cerebral blood volume (δCBV_a_) and total cerebral blood volume (δCBV_tot_). Values of the exponents α_tot_ and α_a_ are calculated on a per subject basis. Group mean and standard deviation weighted by the number of voxels in each subject’s ROI.

Details on how to access the data underpinning the results presented in this study can be found in Appendix A.

## Discussion

A good understanding of the relationship between changes in CBF and CBV is important for interpreting the physiological changes that underlie functional hyperaemia. However, how these changes translate to a measured BOLD response depends on how such changes are distributed across the different vascular compartments. The results of this study will help to improve models of the BOLD response [11,26]. In turn this will enhance the accuracy and reliability of applications that rely on a correct understanding of the BOLD response. For example, it has been shown that the accuracy of the calibrated BOLD method for quantifying stimulus evoked oxygen metabolism changes is critically dependent on accurate knowledge of CBF-CBV_v_ coupling [26,27].

In this study, measurements of CBF, CBV_a_ and CBV_tot_ were combined to assess the coupling of CBF changes with changes in arterial, venous and total CBV, within a CBF-activated ROI. The CBF-CBV_tot_ coupling constant α_tot_ was estimated to be 0.16±0.14 and the CBF-CBV_a_ coupling constant α_a_ was estimated to be 0.65±0.24 (mean±standard deviation). The estimated change in CBV_v_ within the selected ROI was not statistically significant.

### Comparison with the literature

The coupling between CBF and CBV_tot_ has been the target of numerous studies. The power law relationship (Eq. [1]) was first introduced by Grubb *et al*. who measured α_tot_ to be 0.38 in anaesthetised rhesus monkeys during steady state hyper/hypocapnic challenges [5]. Further PET measurements in humans, using a combination of radiolabelled water (H_2_^15^O) and carbon monoxide (C^15^O or ^11^CO), measured α_tot_ to be 0.3 [28] in response to a visual stimulus and 0.29 [29] and 0.64±0.26 [30] during a hyper/hypocapnic challenge (errors reported where available), respectively. MRI based experiments using a perfluorocarbon contrast agent and continuous ASL (CASL) measured a value of α_tot_ of 0.4 in rats [6]. However, in all of these cases measurements were made in the steady state. In this study, a relatively short stimulus duration of 19.2 s was used, which is more relevant for human imaging studies but may not be long enough to allow a steady state to be reached.

The literature regarding the coupling of CBF with CBV_a_ is more limited. Perfluorocarbon studies on rats measured the CBV_a_ change due to steady state hypercapnia and the change in CBF using CASL [6]. Reviewing their data we estimate α_a_=0.84. A similar rat study performed by Kim *et al*. [7] measured changes in CBF and CBV_a_ in response to a 15 s somatosensory stimulus using the MOTIVE technique [31], and from these datasets we estimate α_a_=1.73. The value for α_a_ obtained in our study (0.65±0.24) is at the low end of this range, again perhaps suggesting a steady state was not achieved using a short stimulus. However, further data is required to investigate the effects of stimulus, species, and anaesthesia differences.

This experiment did not allow a direct measurement of CBV_v_ changes. In addition, it was not possible to acquire absolute estimates of CBV_tot_. Therefore the changes in CBV_v_ could only be investigated based on assumptions regarding partitioning of the blood volume. The blood volume was assumed to be described by a two-compartment model consisting of arterial and venous volume fractions. The arterial volume fraction *f*_a_ has previously been measured as 0.27 [7], 0.29 {Duong:2000ia} and 0.3–0.37 [32] using a range of techniques. Each of these methods is expected to include a small fraction of the post-arterial capillary blood volume. The venous volume fraction *f*_v_ has been measured as 0.77 [33] using the quantitative BOLD technique. This technique is specifically sensitive to blood vessels containing deoxygenated blood. Whilst this largely consists of blood within venous vessels, it is also expected to include deoxygenated blood within the capillaries. Therefore, the arterial and venous compartments defined in this study might more accurately be described as the oxygenated and deoxygenated compartments. Importantly, it is this deoxygenated blood volume that underlies the BOLD response and best reflects the BOLD specific CBF-CBV coupling. Therefore, based on the literature values above, *f*_a_ was assumed to be 0.3. Combining Eq. [5] with measurements of ΔCBV_a_ and ΔCBV_tot_, the change in CBV_v_ was predicted to be −0.05±0.11 (Table 1). Across the group, the individual predictions of ΔCBV_v_ were not significantly different from zero. This would suggest a CBF-CBV_v_ coupling constant, α_v_, of around zero. This is low when compared with the limited range of values of CBF-CBV_v_ coupling constant given in the literature. In rats a value of α_v_ of 0.20 was measured using a steady state hypercapnia stimulus [6], whilst in humans α_v_ was measured as 0.18 for a steady state hyper/hypocapnic stimulus [10] and 0.18/0.31 for low/high intensity visual stimulation [9]. However, even in the latter visual stimulus experiments, the stimulus duration was 96 s and therefore significantly longer than that used in our study. This inconsistency between long and short stimuli may be explained by work investigating the temporal dynamics of CBV_v_ [34]. In this work CBV_v_ was shown to be delayed with respect to changes in CBF and CBV_a_. Given the short duration of our stimulus, it is possible that this would not produce an appreciable increase in CBV_v_. Furthermore this is consistent with two photon microscopy measurements that demonstrated a minimal increase in CBV_v_ for a 10 s stimulus, but a larger and delayed response for a 30 s stimulus [35].

Despite this consistency with invasive measurements, we anticipate that the results of this study will be dependent on the ROI selection method. A CBF localiser was chosen to define ROIs in this study to investigate the CBF-CBV coupling properties of the parenchyma, rather than larger venous vessels more distant from the site of activation. CBF ROIs have been shown to be more robust than a BOLD based localiser [36], with the average overlap between CBF and BOLD ROIs of approximately 50%. Combined with the observation that changes in CBV_v_ during activation are spatially heterogenous with both increases and decreases [24], the average CBV_v_ change thus has the potential to vary between different ROI selection methods.

### Limitations of the current study

In this work, ROIs were selected using the activated region defined by the CBF data. However, the use of such functionally defined ROIs can lead to statistical bias, resulting in an overestimation of changes in CBF [37]. Due to the importance of maintaining comfort in human volunteer studies, time within the scanner was limited. Therefore, it was not possible to acquire an additional CBF dataset to provide an independent definition of the ROI.

Absolute quantification of CBV_tot_ was not possible in this study, so we could not measure ΔCBV_v_. This might be measured by tracking the first bolus of contrast agent [38]. However, the resolution of the data acquired in this study was too coarse to yield an adequate arterial input function.

Predictions of α_v_ were based on a two-compartment model of the vasculature: arterial and venous. It was therefore assumed that the capillary compartment was distributed between these two compartments. However, it is likely that changes in capillary CBV will have their own characteristic coupling to CBF and may therefore contribute to an under/overestimation of CBV change in the arterial and venous compartments.

Finally, only steady state changes in haemodynamics were studied since the signal to noise ratio was not sufficient to study dynamic changes. However, it has been shown that a post-stimulus undershoot in CBF may contribute to the BOLD post-stimulus undershoot [39] and that changes in CBV_v_ are delayed with respect to changes in CBV_a_ [34]. These dynamic variations are expected to add to the complexity of the temporal characteristics of the BOLD response. Repeating this work at higher field might allow the study of these dynamic characteristics, greater knowledge of which may enable the dynamics of changes in oxygen metabolism to be investigated using extensions to methods such as calibrated BOLD [1,2].

## Conclusion

In this study, measurements of CBF, CBV_a_ and CBV_tot_ were performed in individual subjects in a single experimental session to assess the coupling of haemodynamic responses. This information is valuable for furthering our understanding of the BOLD response and for enhancing the accuracy and reliability of applications that rely on models of the BOLD response, such as calibrated BOLD.

## Appendix A. Supplementary data

The data that underpin the results presented in this work can be accessed via the Zenodo repository, doi: https://doi.org/10.5281/zenodo.1173621.

## Acknowledgements

This research was funded by the MRC. In addition, RW was supported by a Marie Curie Early Stage Researcher Fellowship and NPB was supported by a Sir Peter Mansfield Postdoctoral Fellowship and by funding from EPSRC grant EP/K025716/1.

